# To Overwrite or to Recall? Individual Differences in Motor Adaptation

**DOI:** 10.1101/439406

**Authors:** Youngmin Oh, Nicolas Schweighofer

**Affiliations:** Neuroscience Graduate Program, University of Southern California, Los Angeles, CA 90089-2520, USA; Biokinesiology and Physical Therapy, University of Southern California, Los Angeles, CA 90089-9006, USA

**Author notes:** Correspondence: Nicolas Schweighofer.

## Abstract

The central nervous system predicts the consequences of motor commands by leaning multiple internal models of external perturbations and of the body. It is not well understood, however, how new internal models are created. Here, we propose a novel computational model of motor adaptation in which a stochastic Bayesian decision-making process determines whether i) a previously updated expert perturbation model is recalled and updated, ii) a novice model is selected and is updated into a new expert, or iii) the “body” model is updated. Results from computer simulations provide insights into various and contradictory experimental data on savings and error-clamp, and predicts qualitative individual differences in adaptation. We verified these predictions in a visuomotor adaptation experiment in which we varied the perturbation amplitudes as well as the amount of noise added to perturbation, and added “trigger” trials in the error-clamp condition. Single trigger trials led to largely qualitatively different behavior and can therefore be used to probe individual differences in memory updates between “one-model” and “two-model” learners. “One-model” learners continuously update the body model, showing no savings during re-adaptation to the perturbation, and gradual decay during error clamp. In contrast, “two-model” learners switch between an updated expert model and the body model, showing large savings during re-adaptation and stochastic lags during error clamp. Our results thus support the view that motor adaptation belongs to the general class of human learning according to which new memories are created when no existing memories can account for discontinuities in sensory data.

**Short summary/significance:** When movements are followed by unexpected outcomes, such as following the introduction of a visuomotor or force field perturbation, or the sudden removal of such perturbations, it is unclear whether the central nervous system updates existing memories or creates new memories. Here, we propose a novel model of adaptation, and investigate, via simulation and behavioral experiments, how the amplitude and schedule of the perturbation, as well as the characteristics of the learner, lead to updates of existing memories or creation of new memories. Our results provide insights into a number of puzzling and contradictory experimental data on savings and error-clamp, as well as large qualitative individual differences in adaptation.

## Introduction

It is now well accepted that the central nervous system (CNS) predicts the consequences of motor commands by leaning multiple models of tools or external perturbations (Wolpert and Kawato, 1998; Krakauer et al., 1999; Lee and Schweighofer, 2009), and by learning internal models of the body (Cothros et al., 2006; Kording et al., 2007; Kluzik et al., 2008). Forward models are updated to minimize sensory prediction errors, i.e., the errors between sensory outcomes and predictions (Mazzoni and Krakauer, 2006; Taylor and Ivry, 2011; Lee et al., 2018). It is thought that if the prediction error is small for one model, then this model will be selected to determine subsequent motor commands and will be further updated if needed (Wolpert and Flanagan, 2001).

It is not well understood, however, how the CNS creates new internal models (Shadmehr and Mussa-Ivaldi, 2012). When movements are followed by unexpected outcomes, such as due to a visuomotor or a force field perturbation, should the CNS update an internal model of the body, should it update one or more perturbation models, or should an entirely new internal model be created?

Here, to address this question, we propose a new model of motor adaptation based on a bank of Kalman filters, along the lines of a recent model of visual memory (Gershman et al., 2014). The model contains three types of internal models involved in motor adaptation: 1) a dedicated body (or “baseline”) model (Berniker and Kording, 2011); 2) “expert” perturbation models that have been previously adapted (Kawato and Wolpert, 1998; Lee and Schweighofer, 2009; Berniker and Kording, 2011); and 3) “novice” unspecific models. As in previous Bayesian models, the decision to select and update a particular internal model when facing a perturbation is based on the models’ prediction errors. If a perturbation is small relative to uncertainty of the baseline model, or if the perturbation is gradually increased, the baseline model is selected and its memory is overwritten. If a relatively large perturbation is re-introduced, or if it is similar to a perturbation encountered in the past, one of the existing expert perturbation models will be selected, leading to the recall of its protected memory. If a new large perturbation is introduced and if it does not match prediction of any existing internal models, then a new novice model is selected and updated. After sufficient training, this novice model becomes a new expert. Besides perturbation amplitude, perturbation noise, motor noise, or individual differences in baseline model uncertainty will influence the effect of the prediction error on model selection and update.

We tested these predictions in a visuomotor adaptation experiment in which we varied the perturbation amplitudes as well as the amount of noise added to perturbation across different experiment groups. At the end of repeating perturbation and washout blocks, a visual error-clamp block was introduced to observe spontaneous and stochastic changes in adaptation. In addition, we inserted two trigger trials (each is a single perturbation trial) in the error-clamp block to test whether a learner has truly developed a new expert model or simply updated the baseline model. The large individual differences in after-effects, savings, and error-clamp can all be accounted for by the uncertainty of the learner in the baseline model.

## Materials and Methods

### Computational model

Our model is based on previous Bayesian models of motor learning known as a mixture of experts (Jordan and Jacobs, 1991; Ghahramani and Wolpert, 1997; Wolpert and Kawato, 1998; Haruno et al., 2001). Specifically, we extend previous models of adaptation based on the Kalman filter (Korenberg and Ghahramani, 2002; Berniker and Kording, 2011) to a model that contains a bank of Kalman filters, along the lines of the model of Gershman et al. (2014) for visual memory. We thus assume that the experts maintain, and udpate, both a mean perturbation estimate and an uncertainty around this estimate. Before being selected for training, these expert models are “novice” with high uncertainty, indicating their non-specificity. In addition, we included a special baseline model (similar to the “body model” in Berniker and Kording (2011)) that estimates the mean perturbation to be zero with low uncertainty and that has a high prior of being selected (Figure 1A).

**Figure 1.**
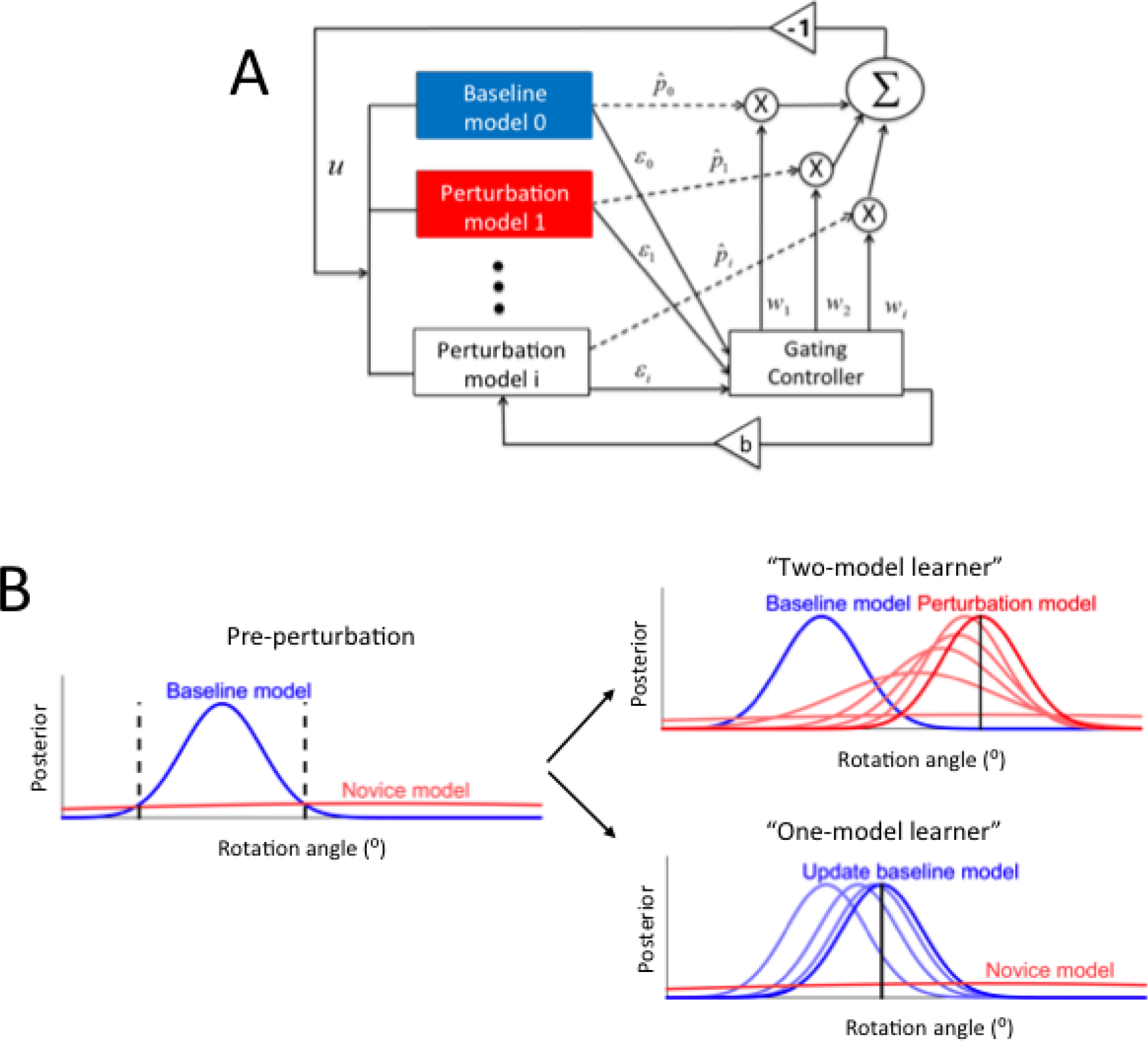
(A) The mixture of experts’ framework. We assumed that the central nervous system maintains a baseline model with relatively low uncertainty and a number of perturbation models. Each model *i* maintains and updates a perturbation mean 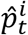 and uncertainty 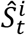. Combination of the model priors and likelihood generates a model weight 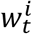 after normalization by the “gating controller”. The weight has two main roles: it controls both the motor output and the model updates. (B) Decision in visuomotor adaptation. Top: because the perturbation amplitude is large with respect to decision boundaries, the learner develops and updates a new expert “Perturbation model”. Bottom: because the perturbation is small, it simply updates the existing “Baseline model”. The light red and blue curves in each panel represent gradual model updates.

As in previous models of visuomotor adaptation, e.g., (Izawa and Shadmehr, 2011), the motor command represents the hand movement direction. On trial *t*, the learner generates a motor command *u*_*t*_ to reach a target *t*_*t*_. Here, we assume that the target is located at the angle 0, that is, the forward direction, without loss of generality. Visual feedback of the hand *h*_*t*_ is determined differently in non error-clamp trials (i.e., baseline, perturbation, and washout) and error-clamp trials, in which feedback is independent of actual performance:

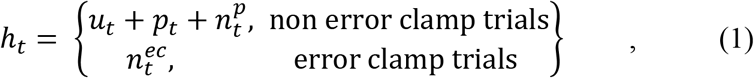

where *p*_*t*_ is the perturbation at time *t*, and 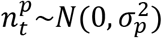 and 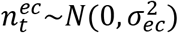 are noise sources added to the perturbation or to the error clamp, respectively.

The estimate of the perturbation 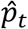 can be given by the weighted predictions from the N models or by the model with the largest weight, in a form of winner-take-all. Although both methods can produce similar results, we chose the winner-take-all approach because it is more robust to changes in parameters. Thus, the overall prediction at time t is given by:

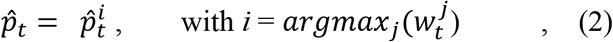

where 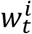 is weight of model *i*, with 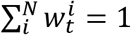.

In order to reach the target at location 0, we assume that subjects generate motor command *u*_*t*_ that compensates the estimated perturbation 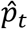:

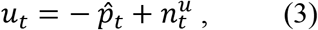

where 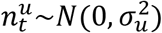 is a motor noise term. Receiving the efferent copy of the motor command, each internal forward model independently predicts the sensory feedback from its own perturbation estimate:

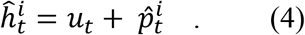

The sensory prediction error for each model is given by:

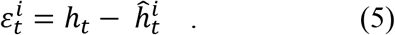

The weights (the “responsibility signal” in previous models such as the MOSAIC models (Wolpert and Kawato, 1998; Haruno et al., 2001; Doya et al., 2002; Bertin et al., 2007)), are given by the posterior probability of the models, given the visual feedback:

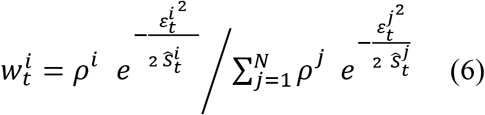

where ρ^*i*^ is a constant prior weight representing prior belief that each model being true in the absence of feedback, and 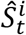 is the uncertainty of a model *i* around its mean estimate 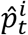. Note that the term in the exponents assumes a Gaussian likelihood function for observations based on predictions.

The predictions and uncertainties are updated accorded to standard Kalman filter equations, e.g., (Bishop and Welch, 2001). However, the type of update for each model in the bank of Kalman filters depends on the weights. When a model is selected, it is updated according to the measurement update equations of the Kalman filter:
for 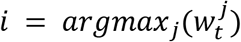:

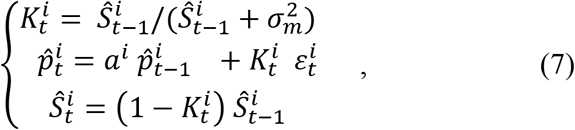

where, 0 <*a*_*i*_ ≤ 1 are decay rate parameters and *σ*_*m*_ a measurement noise parameter. When a model is not selected, it is updated according to the time update equations of the Kalman filter: for *j* ≠ *i*:

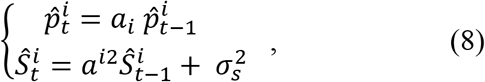

where *σ*_*s*_ a state noise parameter.

### Theoretical predictions

For simplicity, we only consider here a computational model that accounts for adaptation to a single perturbation, with two internal models (Figure 1B): Initially, the model contains a baseline model that estimates the mean perturbation to be zero with low uncertainty, indicating its specificity, and a “novice” model also estimates the mean perturbation to be zero initially but with larger uncertainty, indicating its non-specificity.

The model makes clear predictions that can be tested in simulations and behaviorally. First, it predicts that a large initial perturbation leads to rapid subsequent changes in performance, such as savings and short aftereffects following environmental changes, whereas a small or gradual initial perturbation creates no such rapid changes, but smooth and gradual changes. The former is explained by switching between two expert models while the latter is the result of overwriting the baseline model. More specifically, if an initial perturbation is sufficiently large, i.e., away from the decision boundary defined by the equal weights of the baseline and the novice model, *w*_1_ = *w*_2_ = 0.5 (vertical dashed lines in Figure 1B), the novice model is selected and updated, while the baseline model is protected. After a number of trials, the novice model becomes a new expert: the mean estimate of the perturbation is accurate and the uncertainty becomes small. As a result, the learner can now switch rapidly between these two expert models in subsequent unlearning and relearning blocks, allowing rapid changes of behavior upon environmental changes, expressed as savings in relearning and short or no aftereffects in washout. We call such a learner a “two-model learner”. On the contrary, following a perturbation within the decision boundaries of the baseline model, as would be encountered with a small or gradual perturbation, the baseline model is selected and updated with no update of the novice model. This “one-model learner” will be updated in subsequent washout and relearning blocks of the same amplitude, making transitions slow and gradual, with no savings. In washout, this model will decay gradually back to baseline via trial-by-trial decay.

Second, the model predicts that once a learner has formed two models, switching can occur between the two models in error-clamp. This is because what triggers model switching is the sensory prediction error (equation 6, right panel), but not the performance error, which is clamped to around zero in error-clamp. Thus, assuming that the hand direction is near the adapted direction after a training block, it can show nor or little decay with a lag: the hand direction hovers near the adapted directions until a sudden drop towards baseline. However, high levels of motor noise or experimentally induced perturbation noise increases the probability of yielding a small sensory prediction error for the baseline model, leading to a greater weight for this model, and therefore model switching. Once this happens, performance drops quickly towards baseline. Thus, the model predicts that a sudden drop or rise in performance can occur stochastically in error-clamp, with higher chance with a high level of either perturbation or motor noise. A “rise” can also occur when switching from the baseline to the perturbation model, although this should be a less frequent event than the drop because of the passive memory decay and because of the greater prior assigned to the baseline model.

Next, because the model has qualitatively different modes of operation, single “trigger” trials in the clamp condition can lead to largely qualitatively different behavior when the performance has returned to zero: a trigger trial will cause a sudden rise to the learned level for two-model learners, whereas it will cause only a transient change that decays away quickly for one-model learners.

### Simulations and experimental design

We designed five different sets of simulations to test the above predictions quantitatively. We then performed experiments and data analyses that closely matched the simulation protocols for the first four sets. The simulations and experiments comprised a baseline block, adaptation blocks, washout blocks, and an error-clamp block (Figure 2A).

**Figure 2.**
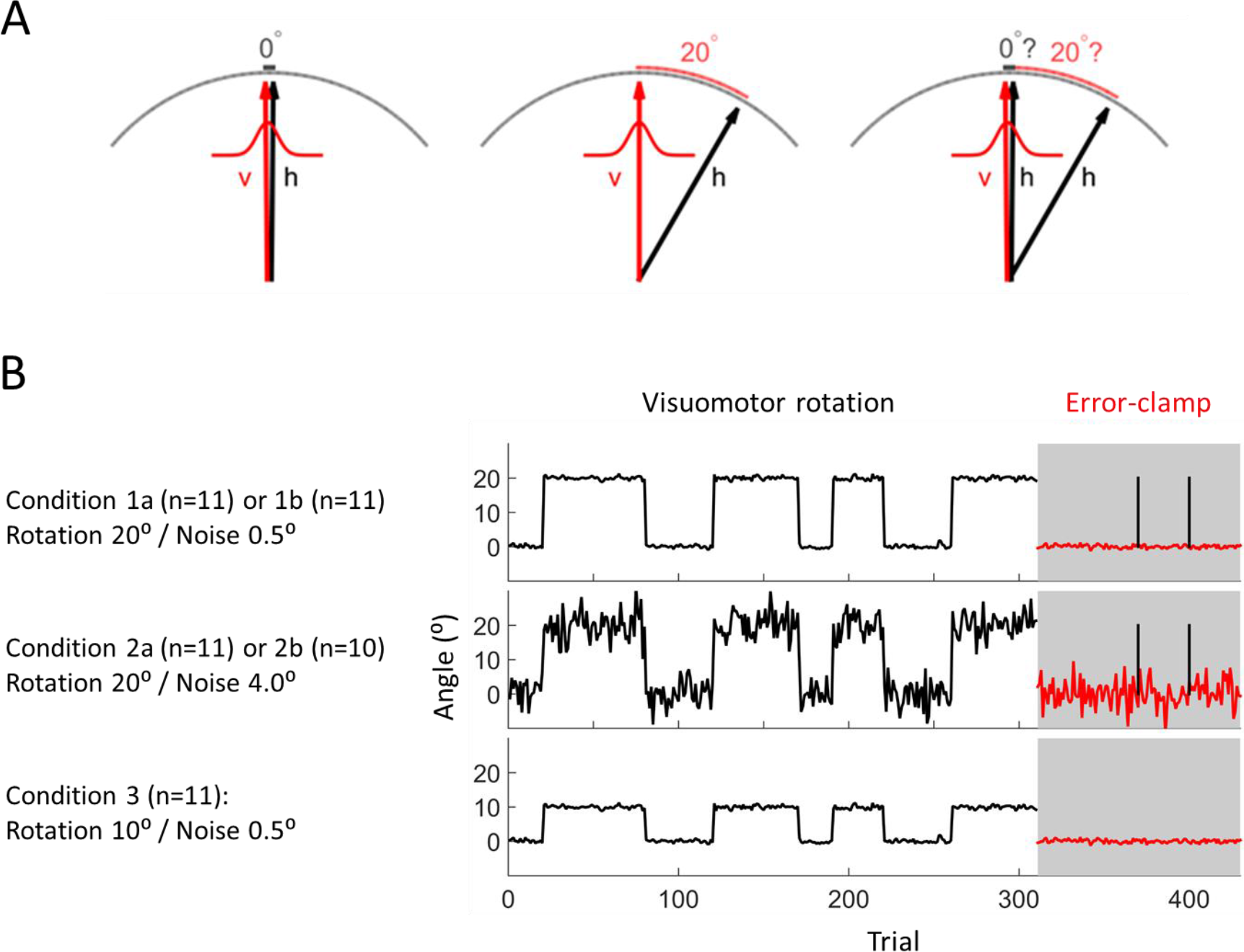
Experiment design. (A) Diagram showing the three stages of the visuomotor rotation paradigm: baseline and washout, adaptation, and error-clamp. Black lines indicate hand movements (occluded to the subjects) and red lines indicate cursor movements. Red curves around the cursor lines indicate the added Gaussian noises to the cursor in experiments. Left panel – baseline and washout: no rotation besides Gaussian noise. Middle panel – visuomotor rotation; the hand movement illustrates behavior at the end of adaptation. Right panel – visual error-clamp: the cursor feedback location is independent of hand direction. (B) The five experimental conditions. Subjects in all conditions experienced the same perturbation schedules except for perturbation magnitude and perturbation noise level. Black lines in adaptation and washout blocks indicate actual rotation, after Gaussian noise added. Red lines in error-clamp indicate actual visual feedback provided. The two black vertical lines in the error clamp show single perturbation (“trigger”) trials given in conditions 1b and 2b.

First, we simulated the effects of large versus small perturbations on subsequent behaviors in unlearning and relearning blocks with a large perturbation (20 degrees) and with a small perturbation (10 degrees). To test for possible time-effect of washout and savings, conditions were simulated with repeated learning and unlearning blocks: baseline (20 trials), learning and unlearning block 1 (60 perturbation trials followed by 40 washout trials), learning and unlearning block 2 (50 perturbation trials followed by 20 washout trials), learning and unlearning block 3 (30 perturbation trials followed by 40 washout trials), and learning block 4 (50 perturbation trials, followed by 120 error clamp trials) (see Figure 2B).

Second, we simulated the effects of large versus small noise levels in error-clamp with large perturbations (20 degrees). The schedule for these simulations was the same as above, but two different noise levels were added to the perturbation. We predicted that the large-noise condition would exhibit smaller lags until decay (i.e., earlier drops from the adapted state to baseline), and the small-noise condition would exhibit longer lags. A density plot of hand directions in the error-clamp block would show distinct distributions for the two conditions: the small-noise condition would have two dominant peaks the distribution of hand directions, one around the learned direction and the other around zero. In contrast, the large-noise condition would have a single dominant peak near zero. The model also predicts that one-model learners, such as found in the small-perturbation condition, would exhibit smooth and intermediate hand distributions between the adapted level and the baseline level in error-clamp, because gradual passive decay occurs in this case, and not model switching.

Third, in addition to the error-clamp simulation described above, we also simulated the effect of two separate trigger trials during error-clamp. The prediction is that if a learner has formed an expert model, then a trigger trial would cause a sudden jump to the learned level if the hand direction were already close to zero before the trigger, and would stay in this up state for several trials following this jump. Otherwise, if the hand were still around the learned level, then a trigger would make no change because it is indistinguishable from the error-clamp trials.

Fourth, to simulate individual differences in adaptation, washout, savings, and error-clamp, we made a change to a single parameter, the baseline model uncertainty (and associated minimum uncertainty 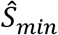), with a first group having a larger uncertainty than the second group.

Finally, we simulated the effects of gradual versus abrupt perturbation, using paradigms akin to those used in Roemmich and Bastian (2015). We tested for savings in a second abrupt adaptation block of 100 trials that is preceded by either a gradual (100 trials), abrupt (100 trials), or short adaptation (20 trials) block.

### Experimental Methods

Fifty-four subjects (22.5±3.8 years old, 20 males and 34 females) participated in the study, which was approved by the Institutional Review Board at the University of Southern California, after signing an informed consent. Subjects sat in front of a device that matched hand space with visual space via a mirror, and were instructed to hold a stylus pen moving on a digitizer tablet (Wacom Intuos 7). Head and trunk movements were limited via a chin-rest. The experiment took place in a dark room, and the mirror-obscured view of the forearm and hand. A cursor (red dot of 1.2 mm radius) representing the tip of the pen was displayed on the mirror. Before the start of each trial, subjects were instructed to position the cursor inside a home circle of a 3 mm radius (about 36 cm away from the subject’s torso). We used a polar coordinate system centered on the home circle, with 0 ° defined as the forward direction and positive direction as clockwise deviation. Subjects were instructed to perform an outward shooting movement toward a circular target of 3 ° radius. The target appeared at a pseudorandom location each trial, within 5 ° around the center of a 120 ° arc that was 10 cm away from the starting position. Subjects were told to initiate a shooting movement as soon as a target appeared and stopped after crossing the arc. A red dot representing a cursor disappeared when the pen tip moved farther than 3 cm from the starting position. When the pen tip crossed the arc, the red dot was displayed on the crossed-point and remained there for 1 s. Subjects were encouraged to keep movement duration between 100 ms and 300 ms, where movement duration was defined as a time interval from the moment when the cursor disappeared to the moment when the cursor crossed the target arc. The messages “Too Slow” or “Too Fast” were displayed when movement duration was out of this range. After each shooting movement, subjects then performed an inward movement to the home circle, during which only the radial location of the cursor was available.

After a familiarization session of 80 trials with no perturbation, the main experimental schedule consisted of repeated learning and washout blocks followed by an error-clamped block. The visuomotor perturbation rotated the cursor position counterclockwise by a given angle with respect to the starting position. The experimental design was closely aligned to the design of the simulations. We randomly assigned subjects into one of five different conditions (Figure 2B): 1a, 2a, 3, 1b, and 2b – see below for a condition description. In all conditions, the experiment schedule consisted of learning blocks was the same order and number of trials as in simulations (see above and Figure 2B). The block lengths were adjusted in pilot tests to allow performance to plateau in each test. The numbers of trials in the learning and unlearning blocks were purposely varied to prevent predictable periodicity in the experiment.

Conditions 1a, 2a, and 3 differed in either rotation angles (Condition 1 vs. Condition 3), or in Gaussian noise levels added to the cursor location (Condition 1 vs. Condition 2). Condition 1a (n=11): large perturbation (20°) and small noise level (std of 0.5°). Condition 2a (n=11): large perturbation (20°) and large noise level (std of 4.0°). Condition 3 (n=11): small perturbation (10°) and small noise level (std of 0.5°). Conditions 1b and 2b were identical to conditions 1a and 2a, respectively, except for two “trigger trials” inserted at half and three quarters of error-clamp, respectively. Trigger trials were simply rotation trials, identical to those in the learning condition (i.e., 20° rotation). In the error-clamp block of all conditions, cursor feedback was independent of actual hand directions, and was sampled from a pre-determined Gaussian distribution of mean 0° and standard deviation 0.5° or 4.0°, depending on the condition noise level. All subjects in each condition received exactly the same rotation sequence.

### Data Analysis

We predicted that the large-perturbation condition exhibits quick changes after the initial learning block whereas the small-perturbation condition is accompanied by slow and gradual changes in all blocks, resulting in large time constants. To test these predictions, we fit performance in each block with an exponential function,

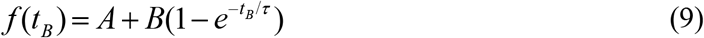

where *t*_*B*_ is a trial number counted from the start of each block, A an amplitude parameter, and τ the time constant. Because the distributions of time constants between groups were not normally distributed, we used Friedman test and Wilcoxon-Mann-Whitney Test to test for differences.

### Simulation parameters

In all simulations, hand direction was modeled between 0 and 1, and then scaled by 20 degrees to match experimental data. We determined a single set of parameters that could replicate all experimental results qualitatively: standard deviations of both sensory noise *σ*_*p*_ and error-clamp noise *σ*_*ec*_ (equation 1) were 0.2 for the large-noise condition and 0.025 for the small-noise condition. The motor noise standard deviation *σ*_*u*_ was 0.15, the measurement noise *σ*_*m*_ was 1 and the state noise *σ*_*s*_ was 0.01. The prior constant of the baseline model ρ^1^ was 0.95 and that of the novice model ρ^2^ was 0.05. Initial mean perturbation values were 0. The initial uncertainties for the baseline and novice (perturbation) model were set to 0.2 and 0.6. In simulations aimed at studying between-subject variability, we increased the initial uncertainty for the perturbation model (and corresponding minimum uncertainty for this model) to 0.35.

Note that to be able to qualitatively reproduce the data, we needed to make two assumptions. First, retention parameters *a* were given different values for the perturbation model and for the baseline and: 0.999 for the perturbation model and 0.98 for the baseline model (See Discussion for justification of this choice). Second, we needed to introduce a minimum value 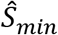 to maintain numeric stability, and took the baseline model uncertainty equal to this minimum value. The minimum uncertainty *S*_min_ was 0.2.

## Results

### Example of simulations illustrating effects of perturbation sizes and noise levels

We first show some examples of simulations for illustrations. In later sections, we present average results over multiple runs. The first column of Figure 3 shows examples of hand directions in the five experimental conditions. The effect of perturbation sizes can be seen by comparing hand direction in the large perturbation conditions (20°; conditions 1a, 2a) to that in the small condition (10°; condition 3; last row). In the large perturbation conditions, hand direction exhibited little aftereffect in washout, large savings in relearning (i.e., one-trial rise), and abrupt switches in error-clamp after varying decay lags or after trigger trials. On the other hand, the small perturbation condition was accompanied by gradual and smooth changes in washout and in relearning blocks, indicating strong aftereffect and no or little savings. Similarly, decay in error-clamp was gradual.

**Figure 3.**
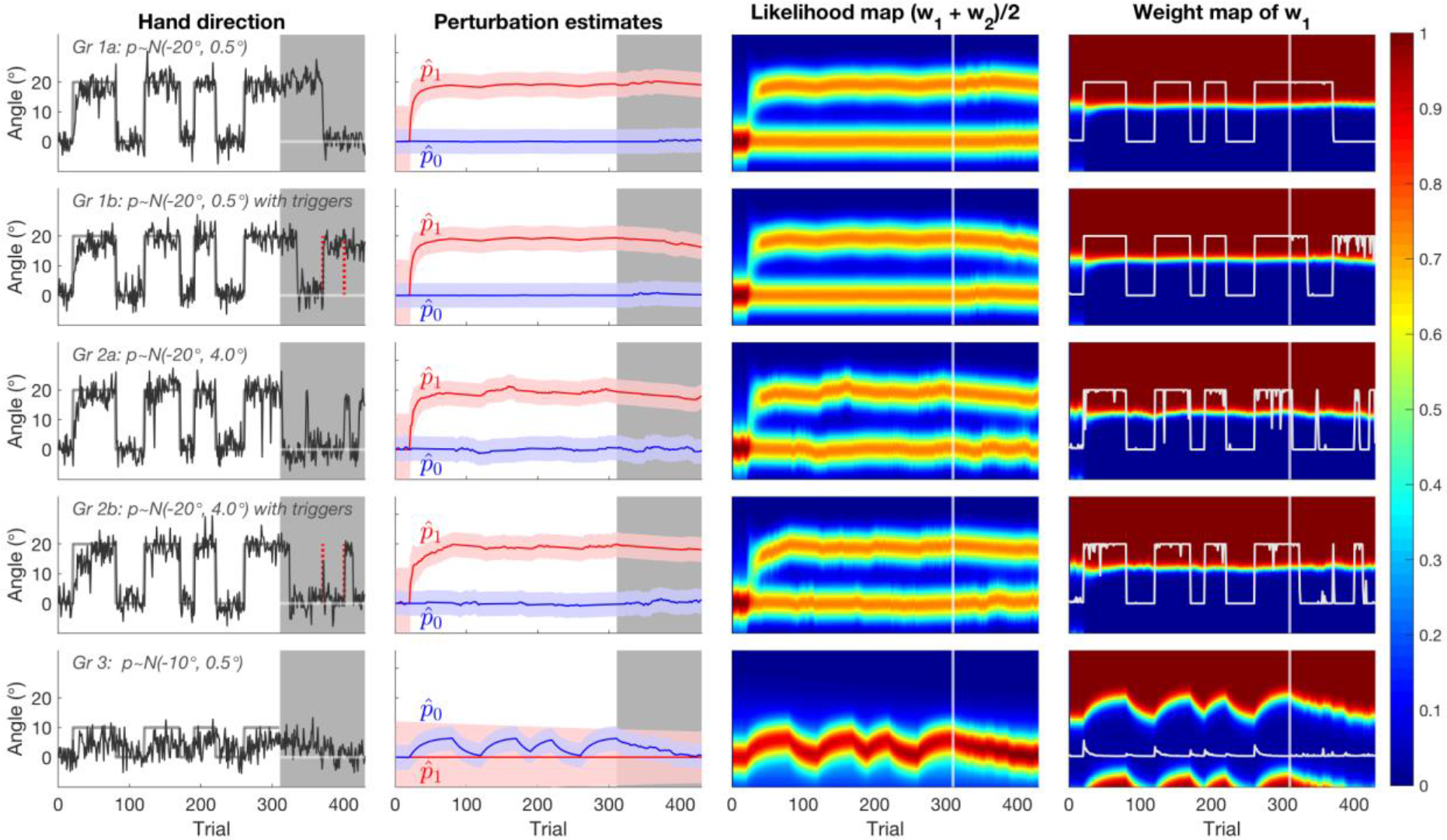
Examples of simulation run for each of the five experimental conditions. First column: black line – negative hand direction, gray line – rotation, gray area – error-clamp block, red dashed line – trigger trial. Second column: perturbation estimation of each learning model. Subscript 0 represents the baseline model (blue), and subscript 1 represents the perturbation model or the novice model if not updated (red). Third column: sum of the two model’s likelihoods (divided by two). Notice how, in the first four conditions, the likelihood of the perturbation model increases as this model better predicts the perturbation, via both the mean prediction convergence to the perturbation level and the decrease in model uncertainty. Fourth column: weight map of the perturbation model, drawn as a function of hand angle and trial. Light gray lines indicate the corresponding weight.

The effect of noise levels can be illustrated by comparing examples of simulations with low levels of noise (condition 1a and 1b) to simulations with high levels of noise (conditions 2a and 2b), in which each pair has identical parameters except for the noise level. Simulations with high levels of noise show earlier switching to baseline in error-clamp than with low levels.

The second column of Figure 3 displays the estimated perturbation mean 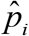, and uncertainty 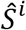 for the baseline and perturbation models. In the large perturbation and small noise conditions (conditions 1a and 1b), the novice model was selected in the initial learning block and its mean estimate increased to approximate the true perturbation level, while its uncertainty simultaneously decreased. The baseline model was practically unchanged. In contrast, in the small perturbation condition (conditions 3), the baseline model updated its estimate of the perturbation up and down each time the environment changed. In contrast, the novice model remained unchanged in this condition, making this learner a one-model learner.

The third column of Figure 3 shows the sum of the likelihoods for both models. As can be seen in the first four conditions, the likelihood of the perturbation model increased as this model better predicted the perturbation. This increase was due to both the mean prediction converging to the perturbation level and the decrease in model uncertainty, as the model was selected (see equation 7).

The fourth column of Figure 3 visualizes the weight map of the perturbation model, superimposed with the weight for this model. As a reminder, because only two models were simulated, the decision boundary is given by *w*_1_ = *w*_2_ = 0.5. In error-clamp, when the perturbation model is near the adapted state, the prediction error is small, and the perturbation model is continuously updated. However, memory decay, perturbation noise, or motor noise, can lead a large prediction error for the perturbation model, yielding a rapid switch to the baseline model. Thus, because the probability of transition was higher for the high perturbation noise level, the lags in error clamps were shorter on average than in the low noise conditions (compare lags in conditions 2a and 2b to those in conditions 1a and 1b). Note that, because of the time-dependent decay of the perturbation estimate, and the higher prior given to the baseline model, the direction of transition is asymmetric, with a higher probability of switching from the perturbation model to the baseline model than in the other direction.

The triggers (vertical red lines in the first columns of conditions 1b and 2b) can act as a cue to signal switching back to the perturbation model when the learner has already returned to the baseline level before a trigger trial. This is shown following the first trigger trials in conditions 1b and 2b. Such switching happens when a single trigger is sufficient to reduce the sensory prediction error for the perturbation model. This transition, however, is probabilistic due to the motor and perturbation noises. Note that if trial-by-trial decay has sufficiently decreased the memory of the perturbation at the time of trigger, there is large prediction error for this model after the trigger, which will therefore be ineffective (results not shown)

### Savings and aftereffects following large and small perturbation: Experimental and simulation results

Figure 4A shows subject-averaged adaptation data for experimental conditions 1a and 1b (20° / 0.5°; top row) combined and for condition 3 (10° / 0.5°; bottom row). Figure 4B shows the mean time constants fitted to the exponential curve (equation 9) within each block across subjects for both learning and washout blocks. Subjects in the large perturbation conditions (20°) showed savings, i.e., significant decrease of time constants in the re-learning blocks (LB2, LB3, and LB4), compared to the initial learning block (LB1; Friedman test; χ^2^ = 34.6, *p* = 1.5 ⟷10^−7^ for conditions1 a/b; χ^2^ = 13.1, *p* = 4.5 ⟷10^−3^ for conditions 2). Post-hoc analysis revealed that savings occurred in all re-learning blocks of conditions 1, with smaller time constants compared to those of the first block (Wilcoxon Signed-Rank Test; *Z* = 3.55, *p* = 4.9 ⟷10^−4^ for LB1-LB2 pair; *Z* = 3.92, *p* = 1.1⟷10^−5^ for LB1-LB3 pair; and *Z* = 3.92, *p* = 1.1⟷10^−5^ for LB1-LB4 pair). On the other hand, no change in time constants of learning blocks was observed in the small perturbation condition (10°; condition 3; Friedman test; χ^2^ = 2.3, *p* = 0.51), indicating a lack of savings. In addition, the overall time constants in washout blocks of conditions 1 were significantly smaller than those in condition 3 (Wilcoxon-Mann-Whitney Test; *Z* = 2.6, *p* = 0.029).

**Figure 4.**
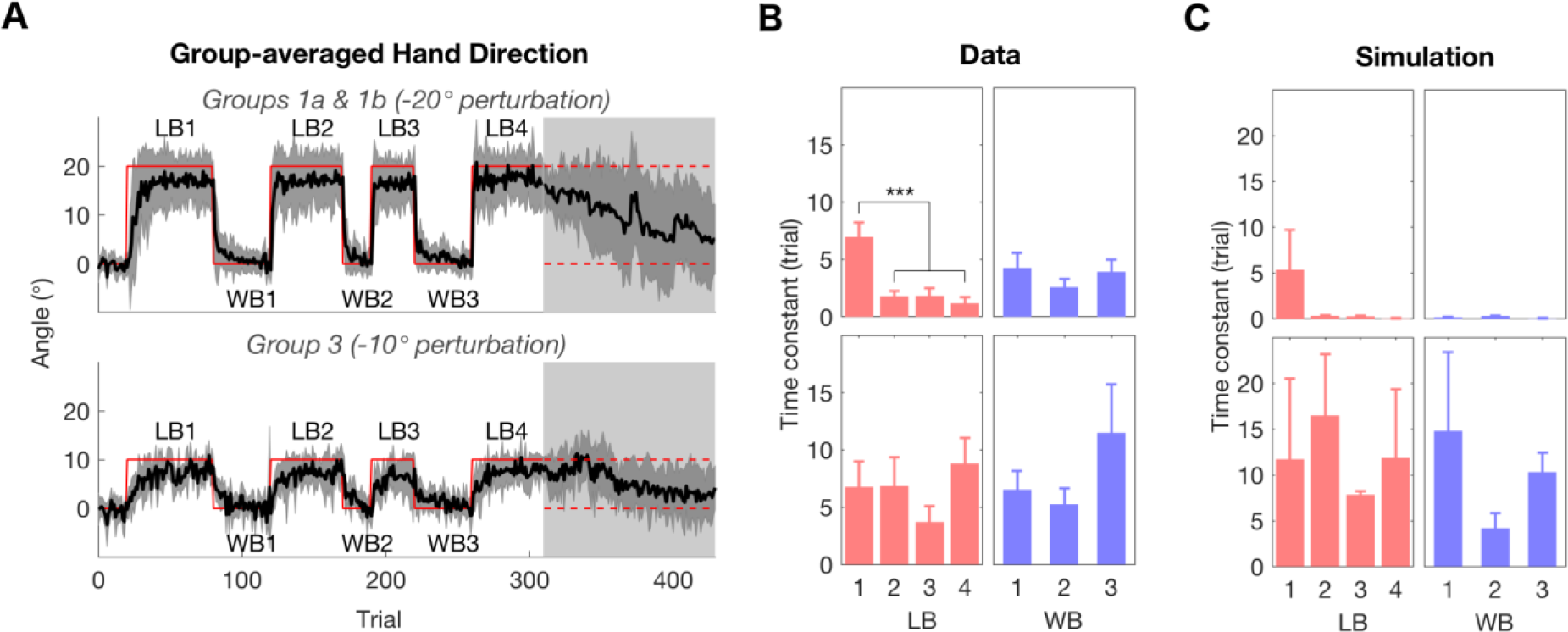
Savings and washout in large and small perturbation conditions. (A-C) Upper row: Conditions 1a and 1b (20° rotation). Bottom row: Condition 3 (10° rotation). (A) Condition-averaged hand direction. Shaded areas around the mean curves (black lines) represent one standard deviation. LB: Learning block (perturbation on), WB: Washout block (perturbation off). (B) Mean time constants for individual exponential fitting from experimental data. Error bars represent standard error. The *** indicate p <0.001 (Bonferroni-corrected). (C) Estimated time constants for each block from simulated data.

In simulations (Figure 4C), the large-perturbation condition produced a median time constant of 16 trials for the first learning block (LB1), followed by small time constants less than 1 trial for the subsequent learning blocks (LB2, LB3, and LB4). The washout blocks produced an averaged median time constants of less than 1 trial across the three washout blocks (WB1, WB2, and WB3). These short time constants were a direct consequence of model switches upon the perturbation change. On the other hand, the small-perturbation condition produced median time constants between 10 and 12 trials for all four learning blocks and median time constants between 11 and 15 for all three washout blocks. These long time constants were a consequence of the baseline model being continuously updated each time the environment changed.

### Decay in error-clamp: Experimental and simulation results

In Figure 5, we show both condition-averaged and individual hand direction in error clamps. Although averaged data suggest a continuous and gradual decay, between-subject variability was large. For instance, for subjects in condition 1a (large perturbation and small noise; Figure 5A), the average between-subject standard deviation of hand direction in learning and unlearning blocks was 3.9°, whereas that in the error-clamp block it was 8.9°. Larger inter-subject variability in error-clamp indicates that dynamics of unlearning in error-clamp may not follow a simple decay. Instead, examination of individual data shows various patterns in error-clamp, with most subjects showing a lag, as predicted by our simulations (see Figure 3). For example, subjects 1, 5, and 10 in condition 1a showed little or no decay, with hand direction above 10° for the whole duration of the clamp. In contrast, subjects 2, 7, and 8 showed a sudden drop after varying lags following the onset of error-clamp. Finally, subjects 4, 6, 9, and 11 showed rather gradual decay. A density plot of hand distribution in error-clamp for all these subjects (Figure 5D, red curve) shows two peaks centered near 0 and 20 degrees. Thus, overall, the hand direction of subjects in condition 1a remained near the perturbation angle of 20° for a relatively large number of trials, and then switched abruptly to near 0°, with few trials between these two angles.

**Figure 5.**
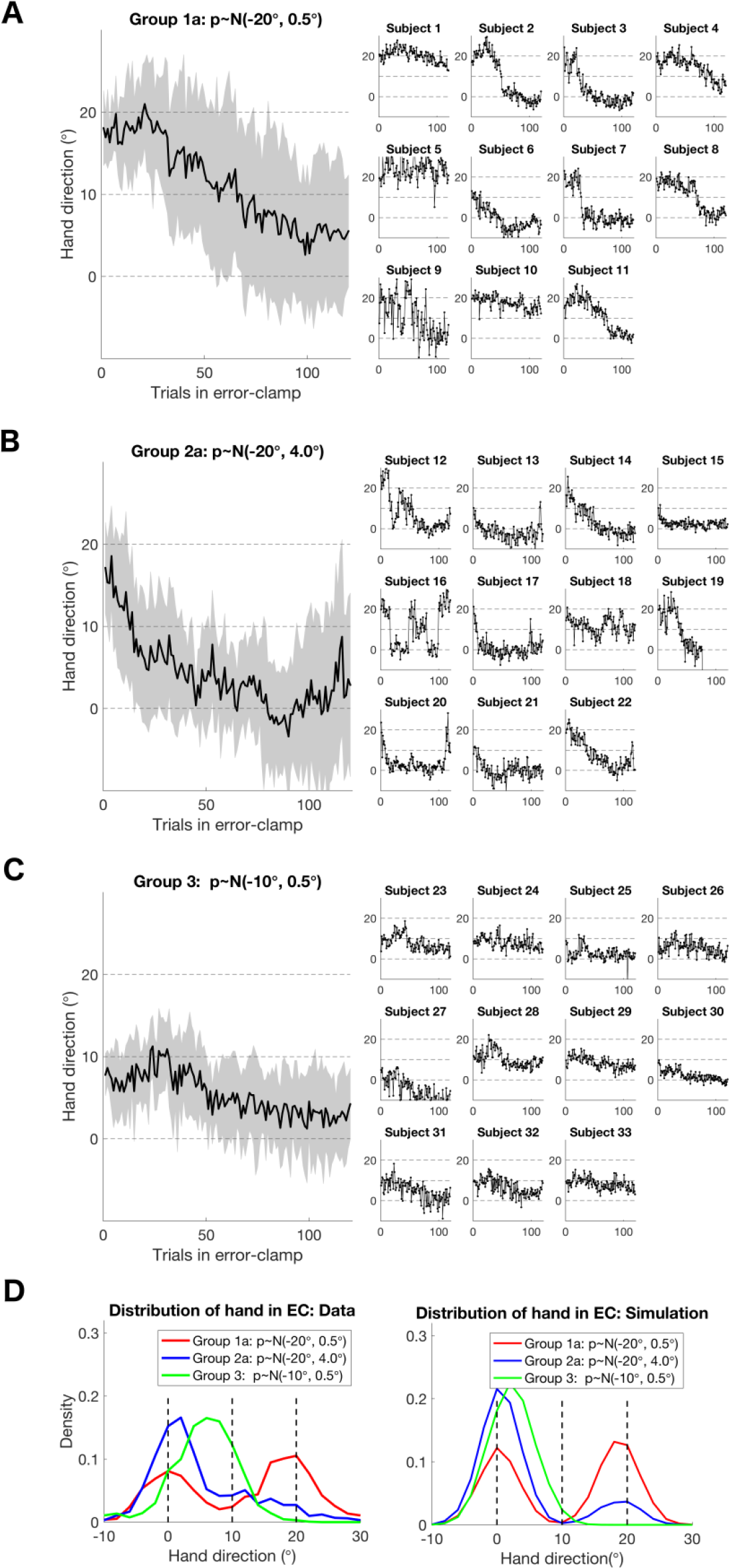
Hand direction in error-clamp. (A) Condition 1a, large perturbation (20°) and small noise (0.5°): Shaded area represents ±1 standard deviation around the condition-averaged plot. (B) Condition 2a, large perturbation (20°) and large noise (4.0°). (C) Condition 3, small perturbation (20°) and small noise (0.5°). (D) Density plot of hand direction in error-clamp for each condition. The left panel shows experimental data, and the right panel shows simulated data.

Average hand direction in condition 2a appears to show faster return to baseline in error-clamp trials compared to condition 1a (Figure 5B), as predicted in simulations. Here again not all subjects followed a simple gradual decay. Subjects 12, 16, 17, 19, and 20 in particular exhibited sudden drops, with subjects 12, 16, and 20 switching back to near 20° spontaneously, resulting in an oscillatory pattern. Note that among those who showed a gradual decay, subjects 13, 15, and 21 started near 10° at the onset of error-clamp; we will discuss a possible cause for this behavior below. A density plot of hand distribution for subjects of condition 2a (Figure 5D, blue curve) shows a single peak centered near 0 degree with a fat right tail. Thus, overall, the hand direction of subjects in this condition also showed sudden switches between perturbation angle and baseline, but such switches occurred earlier than condition 1a with low noise, with occasional spontaneous return back to near 20°.

Condition 3 (small perturbation and small noise; Figure 5C) shows an overall trend of gradual decay and occasional oscillations. The distribution of hand directions suggests that decay was gradual and slow: whereas the density plot in the large perturbation conditions 1a and 2a shows at least one peak near 0°, the distribution of hand direction in condition 3 has a single peak around 8° (Figure 5D, green curve). This suggests that there was no abrupt change, and most subjects did not decay completely to 0° (note however that in this condition, the starting angle in error-clamp was around 10°, and thus trial-by-trial noise made it difficult to distinguish switches, if any, from noise).

For comparison, the right panel of Figure 5D shows the corresponding distributions of hand directions from multiple independent simulation runs of each condition. Compared to data, simulation results show relatively narrower distributions, but they overall replicate the general patterns of density of hand directions in the experimental data, including shapes of distributions and location of peaks in the three conditions.

### Trigger trials and individual differences in adaptation: Experimental and simulation and results

In order to investigate whether a learner had formed two models, i.e., a baseline and a perturbation model, we introduced two trigger trials during error-clamp both in experiments and in simulations of conditions 1b and 2b (Figure 6).

**Figure 6.**
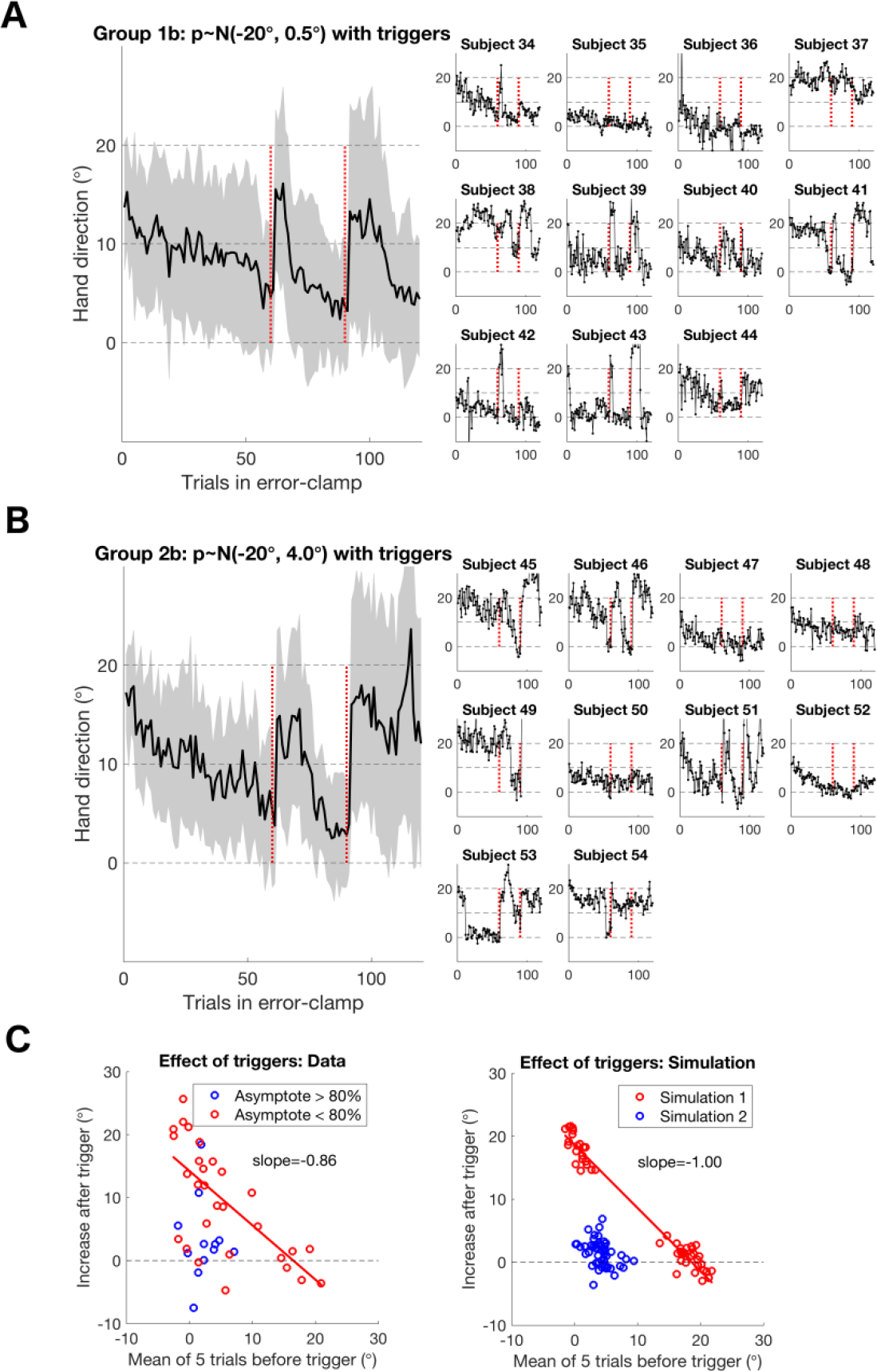
Hand direction in error-clamp with triggers. Red vertical dotted lines indicate a single trigger (perturbation) trial embedded in error-clamp. (A) Condition 1b, large perturbation (20°) and small noise (0.5°). Shaded area represents ±1 standard deviation around the condition-averaged plot. (B) Condition 2b, large perturbation (20°) and large noise (4.0°). (C) Effect of the triggers for the 21 subjects of groups 1b & 2b. The X-axis represents the mean hand direction of 5 trials before each trigger trial, and the Y-axis represents the change in mean hand direction (i.e., mean of 5 trials after trigger minus mean of 5 trials before trigger). Left panel (Data) – each circle represents a single trigger event; since there are 2 triggers for each subject, there are 42 events (circles). The red circles indicate trigger events from subjects who adapted to more than 80 % of perturbation in the last 10 trials of the last block. The blue circles indicate the other subjects. Right panel (Simulation) – each small circle represents a simulated trigger event from 50 independent simulation runs. Red circles represent two-model learners, who developed two internal models, whereas blue circles represent one-model learners, who only updated the baseline internal model.

Condition-averaged hand direction in error-clamp showed instantaneous responses to the triggers (i.e., sudden jumps in hand direction), both in conditions 1b and 2b. The response appears sustained for a number of error-clamp trials thereafter (Figure 6A and 6B, left panels). However, the averaged hand direction indicates that the responses to triggers were, on average incomplete in magnitude, i.e., smaller than 20°. This is because not all individuals responded to the trigger; indeed, three patterns can be observed. First, when the hand direction was near 0° when a trigger was presented, a majority of subjects (subjects 34, 38, 39, 41, 42, 43, and 44 of condition 1b; 45, 46, 49, 51, 53 and 54 of condition 2b) showed immediate jumps in response to the trigger to angle values near the adapted state. In contrast, and as expected when the hand direction was near 20° at the time of trigger, there was no effect (see subject 37, 38 of condition 1b and subject 45, 49, and 54 of condition 2b). These first two patterns were predicted by our model – compare results for these subjects with the model’s response to trigger in Figure 3A.

However, a third pattern is also apparent: subjects with hand direction near 0° at the time of trigger who did not respond to the trigger (subjects 35 and 36 of condition 1b; subjects 47, 48, 50, and 52 of condition 2b). We note that those non-responding subjects appeared to have started the error-clamp block with a lower level of adaptation than those who responded. We therefore hypothesized that subjects who fully adapted to the perturbation formed a new perturbation model, that is, were two-model learners. In contrast, subjects who did not fully adapted to the perturbation only updated the baseline model, that is, were one-model learners.

To test this hypothesis, we divided subjects in conditions 1b and 2b between “full-learners”, as defined by subjects whose asymptotic adaptation angle in the last learning block (LB4) was greater than or equal to 80% of the full angle (so more or equal to 16 °), and “partial-learners” if below 16 °. Figure 6C (left panel) shows a strong negative correlation between the hand direction before the trigger trials and the amount of hand change after the trigger trials for the full-learners. The slope of −0.86, which is close to −1, indicates that the hand direction switched to around 20° after the trigger, independently of hand direction before the trigger. In contrast, the partial-learners failed to show such a relationship. As can be seen by the cluster of blue circles on the bottom left of the figure, most subjects who had a lower adaptation level at the end of adaptation failed to respond to the trigger trials.

Figure 6C (right panel) shows simulation results that account for these patterns of response to triggers for 20 simulated subjects. Two sub-groups of subjects were simulated: a sub-group with default baseline uncertainty (and with the associated minimum uncertainty) of 0.2, the two-model learners; and a sub-group with a broader baseline uncertainty of 0.35, the one-model learners (Note that because the model parameters are scaled to a maximum perturbation of 1, the two-model and one-model learners baseline model uncertainty correspond to 4° and 7°, respectively). The sub-group with the default, more narrow, baseline uncertainty developed two models during adaptation, and responded to triggers in clamp, as shown in Figure 3. In contrast, the sub-group with the broader baseline model uncertainty did not develop a new perturbation model, and therefore behaved as the one model learner in the last row of Figure 3 (but of course, trying to adapt to a 20° instead of a 10° perturbation). Thus, in our paradigm, 20° appears to be a large perturbation for most subjects, leading to the development of a new model. For several other subjects however, the same 20° perturbation appears to only warrant the update of the baseline model.

Based on the savings results of large and small perturbation shown in Figure 4, we then conjectured that the subjects who showed low levels of adaptation at the end of the last adaptation block, and therefore who, for the most part, failed to respond to triggers, would show little or no savings in relearning blocks as well as gradual after effects in washout blocks. We therefore performed an analysis similar to the savings analysis in Figure 4, and calculated the time constants for the washout and learning blocks for the “full learners” and “partial learners” sub-groups (as defined by the 80% threshold discussed above) for all the four 20° perturbation conditions (groups 1a, 1b, 2a, and 2b). This resulted in 33 full learners and 10 partial learners. Figure 7 shows that indeed, the full-learners showed large savings and short after-effects, whereas the partial-learners showed little savings and strong after-effects. The overall time constants of relearning blocks (mean of LB2, LB3, and LB4) of the full-learners were significantly smaller than those of the partial-learners (Wilcoxon-Mann-Whitney Test; *W* = 530, *p* = 0.0001), despite the fact that their initial learning was not significantly different (comparing LB1; *W* = 358, *p* = 0.62). In addition, the overall time constants of washout blocks (mean of WB1, WB2, and WB3) of the full-learners were significantly smaller than those of the partial-learners (*W* = 542, *p* = 0.00005).

**Figure 7.**
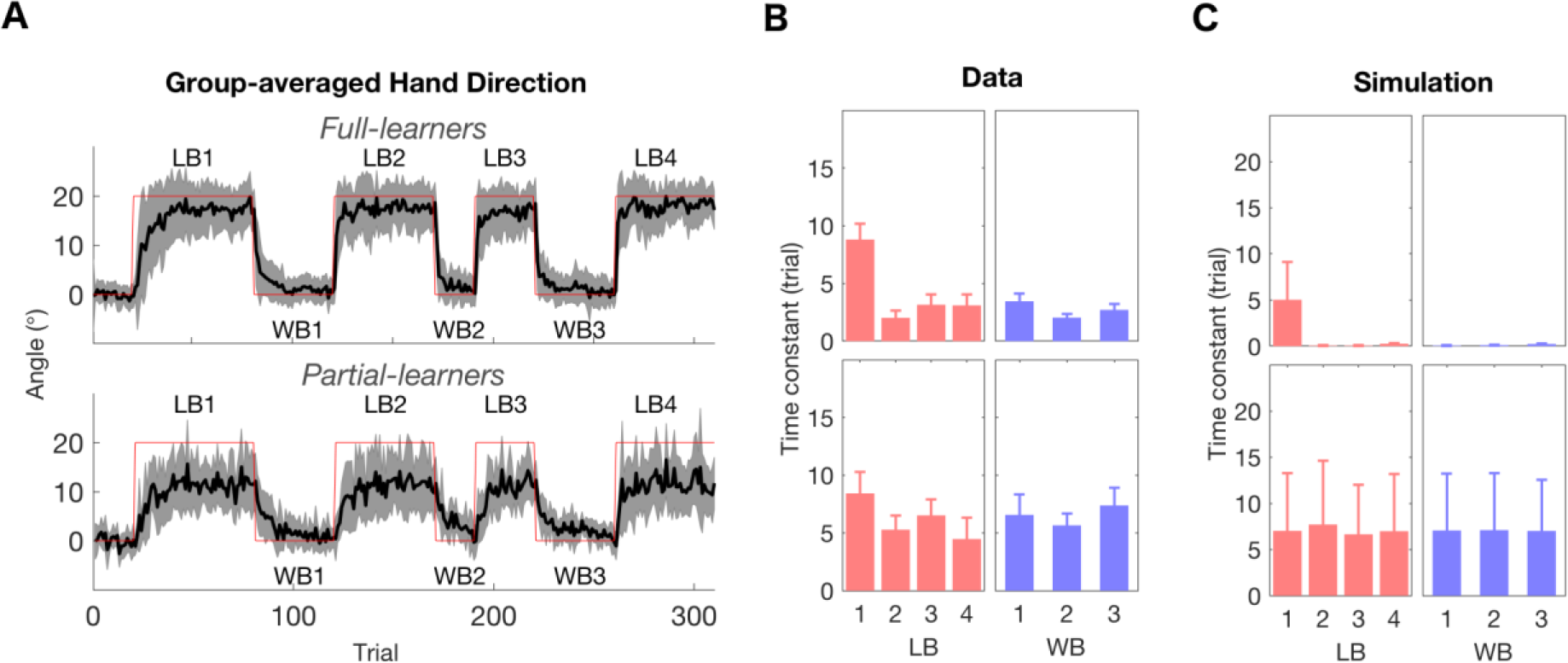
Savings and washout for Full-learners vs. Partial-learners. (A-B) Upper row: Full-learners (Asymptote of LB4 >= 16°, N=33). Partial-learners (Asymptote of LB4 < 16°, N=10) Condition-averaged hand direction. Shaded areas around the mean curves (black lines) represent one standard deviation. LB: Learning block (perturbation on), WB: Washout block (perturbation off). Mean time constants for individual exponential fitting from experimental data and simulation data (C). Error bars represent standard errors.

### Savings and aftereffects following gradual and abrupt perturbations: Simulation results

Here we present simulations that account for “one-trial” savings in the data from Roemmich and Bastian (2015). Specifically, these authors showed that whereas a gradual perturbation followed by a washout period does not lead to savings in a subsequent abrupt re-adaptation phase (their phase G1WA2), an abrupt perturbation followed by a washout period leads to large savings in a subsequent abrupt re-adaptation phase (AWA2). A short abrupt perturbation followed by a washout period also leads to savings in a subsequent abrupt re-adaptation phase (sAWA2). Figure 8A (left) shows the hand direction in the three conditions for a single simulation runs. Figure 8B shows the average of 50 simulations, in which, similar to Figure 4 in Roemmich and Bastian (2015), we superimposed the adaptation A2 in G1WA2 to the initial abrupt adaptation A in AWA2 (top), the adaptation A2 in AWA2 to the initial abrupt adaptation A in AWA2 (middle), and the adaptation A2 in sAWA2 to the initial abrupt adaptation A in AWA2 bottom. As in previous experiments in locomotion (Roemmich and Bastian, 2015) and arm movements (Herzfeld et al., 2014), the gradual adaptation GWA2 resulted in no savings. In contrast, the initial abrupt adaptation in AWA2 resulted in large “one-trial” savings and, to a lesser extent, the short initial abrupt adaptation in AWA2 also resulted in savings.

**Figure 8.**
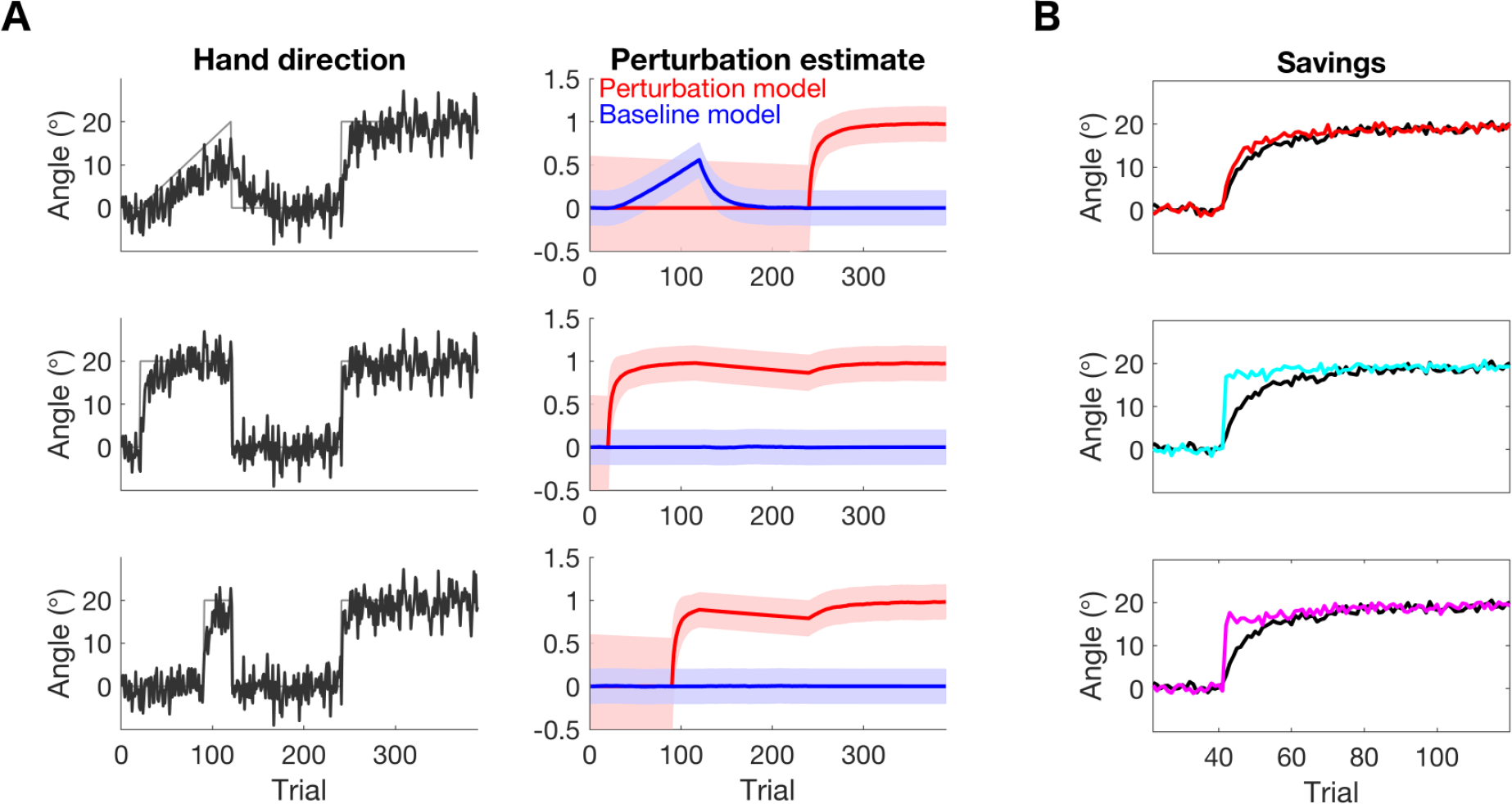
Simulated savings following gradual (top), abrupt (middle), and short-abrupt (bottom) initial perturbations. A. Single runs illustrating hand directions (Left) and model updates (middle) in the three conditions. B. Average of 50 simulations, in which the initial and second perturbations are aligned at trial 0. Note how savings are very strong following a long abrupt perturbation, still important following a short-abrupt perturbation, but there are no savings following the gradual perturbation. Compare with Figure 4 in Roemmich and Bastian (2015).

## Discussion

The significance of our study is that we propose a unifying model of motor adaptation that accounts for a variety of phenomena that appear, *a priori*, contradictory. Our key novel insight is that qualitative differences in learning, washout, savings, and decay in error-clamp between conditions and between participants are all linked and explained by a single computational model that uses a stochastic Bayesian decision-making process. At every trial, the sensory prediction error of the internal models will determine whether i) a previously updated expert model is recalled and updated, ii) a novice model is selected and develop into a new expert, or iii) the baseline model is updated. Because of these three different modes of operations, the model generates qualitative differences in savings and during error clamps.

### Savings following large vs. small perturbations and abrupt vs. gradual perturbations

As predicted by the model, experimental data revealed that savings were dominant in the large, but not in the small, perturbation conditions. In contrast, long-lasting after-effects were observed in the small, but not in the large, perturbation condition. These results partially replicate those of Morehead et al. (2015), with the main difference that we used 20° and 10° for the large and small perturbation with a single target, whereas they used 45° and 15° with two targets. In addition, our results are in line with those of previous studies showing that savings in motor adaption depend on the condition of the perturbation. On one hand, savings occur after an initial large abrupt perturbation followed by a washout period (Klassen et al., 2005; Krakauer et al., 2005; Morehead et al., 2015), even with very few trials of adaptation (Huberdeau et al., 2015), and after a gradual perturbation if followed by a no-activity period (Klassen et al., 2005). On the other hand, no savings occur when a gradual adaptation is followed by a washout period (Herzfeld et al., 2014; Roemmich and Bastian, 2015), unless there was a large error at the end of the gradual adaptation block (Turnham et al., 2012).

Previous models cannot account for these various savings results. Linear time-invariant models can only account for savings following very short washout sessions (Smith et al., 2006). Models with multiple memories can account for savings following abrupt perturbations because they protect existing memories. However, the Lee and Schweighofer (2009) model needed an external contextual cue for long-term update, and thus cannot account for savings after gradual adaptation, and the Berniker and Kording (2011) model predicted savings following gradual adaptation, unlike the data of Roemmich and Bastian (2015). In contrast, our current model explains these results because only the baseline model is updated during gradual adaptation.

### Stochastic lags versus gradual decay in error-clamp

Our model and experimental results allow reconciliation of previously controversial experimental results on error-clamps. It has been shown that performance during clamp can be sustained near the perturbation level and decay stochastically after varying lag durations (Scheidt et al., 2000; Vaswani and Shadmehr, 2013; Vaswani et al., 2015). In stark contrast, Brennan and Smith (2015) proposed that the lag observed in the Vaswani and Shadmehr (2013) study was an artifact, and that the decay in error-clamp was due to trial-by-trial passive forgetting. Our data show both lags and gradual decay, with the conditions of adaptation controlling the decay behavior in the clamp. Large perturbation amplitudes and low noise levels promote long stochastic lags followed by abrupt decay. Large amplitude and large noise levels promote shorter lags. Small amplitude or gradual adaptation promotes gradual decay. In addition, following large perturbations, subjects showing higher adaptation levels exhibit lags, whereas subjects showing lower levels exhibit gradual decay.

The distinction between two-model learners and one-model learners can reconcile these error-clamp data. If the learner becomes a two-model learner, then lags and model switching based on prediction error occurs. Contrarily, if the learner only updates the baseline model, then passive gradual decay occurs. In addition, increase in motor or sensory noise leads to greater noise in sensory prediction error, and therefore increasing chance of switching to the baseline model, with shorter lags. To demonstrate that switching can occur in error-clamp, we inserted trigger trials in the error-clamp following adaptation to large perturbation. The data often showed fast switches and sustained performance just after these single trials. Some subjects did not respond to trigger trials. However, these subjects were partial learners, which suggests that they only updated their baseline models.

Recent research (Taylor et al., 2014) revealed that adaptation has both a “strategic” and implicit components. Strategic learning is temporally-stable, while implicit learning consists of both temporally-stable and temporally-labile components (Miyamoto et al., 2014). Involvement of such strategic learning could explain why, in our simulations, a larger decay rate was needed in the baseline model than in the expert models to account for decay in the clamp data (large decay rate in the expert models would have created much shorter lags). This suggests that update of a newly learned model involves both strategic (temporally-stable) and implicit (both stable and labile) learning, whereas update of the baseline model mostly involves implicit learning.

### Limitations and future work

A first limitation of our model is that it accounts for only one of the two types of savings (Roemmich and Bastian, 2015), the initial savings shown in the first trials of re-adaptation. Later savings, seen by an increase in learning rate require a meta-learning process, e.g., (Schweighofer and Doya, 2003; Herzfeld et al., 2014). Future models will need to incorporate these two types of savings to provide a full account of existing experimental data. A second limitation is that we assumed, for the sake of simplicity, that only prediction errors update motor adaptation. However, motor adaptation is also updated by reward-based and use-dependent mechanisms, e.g., (Huang et al., 2011; Shmuelof et al., 2012; Galea et al., 2015). Because such additional mechanism influence savings (Huang et al., 2011), dissociation of the multiple different factors that influence savings is needed. A third limitation is that we did not model processes with different timescales involved in motor adaptation. Previous studies have shown the existence of fast and slow processes (related to the temporally stable and labile processes discussed above)(Smith et al., 2006), possibly updated in parallel (Lee and Schweighofer, 2009); recent studies suggest that additional time processes exist (Kording et al., 2007; Kim et al., 2015). Future work will be needed to understand how to integrate these fast/slow processes with the multiple memories framework discussed in this paper.

### Related models

Our main question was to study how the CNS creates new internal models (Shadmehr and Mussa-Ivaldi, 2012). The question of updating existing models been addressed via the mixture of experts’ models, e.g., (Jordan and Jacobs, 1991; Ghahramani and Wolpert, 1997; Wolpert and Kawato, 1998; Haruno et al., 2001; Lee and Schweighofer, 2009; Schweighofer et al., 2011; Lee et al., 2016). While these previous models used experts to explain adaptation to multiple tasks, the current model and a previous model (Berniker and Kording, 2011) suggest that modular experts are needed to explain both learning and forgetting of a single task.

In addition, whereas these previous models of adaptation assumed the existence of a finite set of experts a priori, we assumed here the existence of novice models. Similarly, Lonini et al. (2009) proposed that new local regression models are added whenever a new data point does not activate any of the existing models above a threshold. As in our model, previously learned memories are protected, thus the model can show savings. However, it is unclear, how their model could account for data with small or gradual adaptation and for the qualitative difference in error-clamps.

Our model is highly related to a previous model of visual memory (Gershman et al., 2014). Like our model of motor adaptation, this model is based on a bank of Kalman filters, and new memories are created when discontinuities in sensory data cannot be explained by existing memories. Thus, it is striking that a similar mechanism, based on prediction errors, may be involved in the formation and update of different types of memories (visual memory and motor memory) in presumably different brain areas, with the cerebellum involved in visuomotor memories, e.g., (Kim et al., 2015). Thus, our simulation and behavioral data of motor adaptation are in line with the general view of human learning according to which new memories are created when no existing memories can account for discontinuities in sensory data (Gershman et al., 2014).

In these models, the width of the “receptive field” of the prediction error determines if the new data is sufficiently “close” to the memory. We proposed that the width of the baseline model is subject-specific. Accordingly, what constitute a large or a small perturbation is variable between subjects and may depend on the uncertainty of the baseline model being within “natural error range” (Torres-Oviedo and Bastian, 2012). Additional studies are needed to determine the neural correlates underlying one-model versus two-model learning in motor adaptation.

## Acknowledgements

We thank Jun Izawa for helpful discussions on models of motor adaptation and Raphael Schween for valuable comments on a previous draft. This work was funded by grants NSF BCS 1031899, NIH R01 HD065438 and R56 NS100528 to NS.

